# Epilepsy-associated potassium channel KCNT1 is required for multiciliated cell development in Xenopus

**DOI:** 10.64898/2026.03.31.710877

**Authors:** Angeline K. Chemel, Kate E. McCluskey, Matthew N. Tran, Aliza T. Ehrlich, Helen Rankin Willsey

## Abstract

Pathogenic variants in the gene *KCNT1*, which encodes a sodium-activated potassium channel, cause a severe neurodevelopmental disorder with intractable epilepsy. In addition to seizures, affected individuals commonly present with severe respiratory issues and structural heart defects not commonly observed in other genetic pediatric epilepsies, suggesting additional developmental functions for KCNT1 in organs beyond the brain. Here, we characterized the spectrum of clinical diagnoses present in a cohort of 46 individuals with pathogenic variants in *KCNT1*, ranging from 0 to 19 years of age, by medical record review. We documented the prevalence of diagnoses across organ systems, including dependence on assisted breathing, congenital structural heart defects, urinary dysfunction, and spine deformities, among others. Next, we explored the embryonic expression and function of KCNT1 in diploid frogs (*Xenopus tropicalis)* and observed expression in developing ciliated tissues such as the brain, heart, kidney, and epidermis. Embryonic perturbation of KCNT1 disrupted developmental signaling pathways and caused ciliogenesis defects in the mucociliary epidermis, a common model for the human airway. Loss of KCNT1 disrupted development of multiciliated cells, reminiscent of recent work on the ion channel Piezo1. Consistently, pharmacological inhibition of Piezo signaling enhanced the ciliogenesis phenotype observed following KCNT1 inhibition, while activation of Piezo1 activity partially rescued ciliogenesis in the context of KCNT1 inhibition. Together, this work establishes that KCNT1 has embryonic functions in *Xenopus* beyond regulating neuronal activity, specifically in multiciliated cell development, and identifies an interaction with pharmacologically-tractable Piezo channels that may be productive for therapeutic efforts.

## Introduction

Pathogenic genetic variants in *KCNT1*, potassium sodium-activated channel subfamily T member 1, cause early-onset epileptic encephalopathies (Barcia et al., 2012), marked by profound developmental delay, intellectual disability, and significant impairment in breathing, feeding, and walking (Bonardi et al., 2021). Disease-associated *KCNT1* variants are almost exclusively missense mutations predicted to confer gain-of-function activity and cluster in the C-terminal region near the potassium conductance domains (Barcia et al., 2012; Hite et al., 2015). Clinically, KCNT1-related epilepsy is highly pharmacoresistent, with most antiseizure treatments producing little to no meaningful reduction in seizure burden and highly variable responses across patients (Fitzgerald et al., 2019; Gras et al., 2024; Mikati et al., 2015).

What distinguishes KCNT1-related disorders from many other genetic intractable pediatric epilepsies is the striking breadth and severity of systemic involvement beginning early in life (McTague et al., 2018). In addition to catastrophic epilepsy, affected children frequently exhibit life-threatening comorbidities, including severe respiratory dysfunction requiring chronic ventilatory support (Bonardi et al., 2021), which is uncommon in most monogenic epilepsies. Mortality is correspondingly high, with deaths often attributed to sudden unexpected death in epilepsy (SUDEP) and respiratory failure (Kuchenbuch et al., 2019). Even more unusual is the recurrent presence of structural cardiovascular abnormalities, including systemic-to-pulmonary collateral vessels (Delaney et al., 2025) and other aberrant vascular connections originating from the heart that can lead to hemoptysis and heart failure (Kawasaki et al., 2017; Kohli et al., 2020). These extracerebral manifestations are sufficiently prevalent that recent work has proposed expanding the clinical spectrum of KCNT1-related disease to explicitly include cardiovascular and vascular defects (Kuchenbuch et al., 2019). Together, this constellation of early, multi-organ symptomatology points to KCNT1-related epilepsy as not solely a disorder of neuronal excitability, but a broader developmental channelopathy with systemic consequences. This perspective raises critical questions about when and where KCNT1 functions during development and suggests that effective therapies may need to address pathogenic mechanisms that extend well beyond seizure control. Despite insights from KCNT1-deficient mouse models into behavioral phenotypes such as learning and memory (Bausch et al., 2015), the endogenous developmental role of KCNT1 remains poorly defined. In particular, its embryonic expression pattern has not been systematically characterized, and the developmental consequences of KCNT1 dysfunction across organ systems remain unknown. Addressing these gaps is essential for understanding how *KCNT1* mutations produce such an unusually severe and multisystem disorder.

*Xenopus* provides a powerful *in vivo* platform for interrogating embryonic gene function and has been instrumental in uncovering conserved mechanisms governing airway and heart development and disease (Duncan and Khokha, 2016; Hempel and Kühl, 2016; Rankin et al., 2015; Walentek, 2021; Walentek and Quigley, 2017; Warkman and Krieg, 2007). Notably, in *Xenopus*, potassium channels have been shown to regulate key embryonic processes, including gastrulation and left-right patterning (Sempou et al., 2022), highlighting the emerging importance of ion flux and membrane potential in embryonic morphogenesis. Cilia, membrane-bound cellular organelles, sit at the intersection of several phenotypes prominent in KCNT1-related disease, as cilia defects can cause intractable epilepsy, respiratory dysfunction, and congenital heart defects (Reiter and Leroux, 2017). Further, the cilia-dependent hedgehog signaling pathway regulates pulmonary vessel quiescence (Peng et al., 2015), raising the possibility that ciliary dysfunction may be relevant to KCNT1-related disorders. Consistently, membrane potential and ion channel activity are increasingly recognized as regulators of ciliogenesis (Alshriem et al., 2025; Kulkarni et al., 2021; Narayanan et al., 2025; Saternos et al., 2020; Ventrella et al., 2023), and an epilepsy-associated potassium channel, *KCNH1*, has recently been shown to regulate cilia development (Napoli et al., 2022; Sánchez et al., 2016). These observations led us to hypothesize that KCNT1 may have an additional unrecognized role in ciliary biology that contributes to the multisystem features of the disorder.

Here we combined clinical phenotyping with developmental *in vivo* modeling to test this hypothesis. Through medical record review of 46 individuals with pathogenic *KCNT1* variants, we quantified the prevalence of respiratory, cardiac, musculoskeletal, renal, and dysmorphic features. We then mapped *KCNT1* expression during *Xenopus* embryogenesis and observed strong expression in developing, highly ciliated tissues. Using both genetic and pharmacologic approaches to model KCNT1 loss-of-function, we uncovered a requirement for KCNT1 in multiciliated cell development. Mechanistically, we identify an interaction with Piezo signaling, consistent with prior links between potassium flux and Piezo activation (Mitchell et al., 2025) and between the cilia specification regulator FOXJ1 and Piezo activity (Narayanan et al., 2025). Together, our findings reveal a previously unappreciated embryonic role for KCNT1 in ciliary biology and provide a developmental framework for understanding the unusually broad clinical spectrum of KCNT1-related disease. These results also suggest new avenues for therapeutic targeting of Piezo1-mediated pathways in this devastating disorder.

## Results

### Medical record review of individuals with KCNT1 variants

To formally document the range of symptoms in individuals with *KCNT1* variants, we reviewed patient medical record data provided by Citizen Health. These data were compiled from 46 patients with pathogenic variants in *KCNT1*, ranging from 0 to 19 years of age, 44% female and 56% male. We documented the rates of seizures (100%), respiratory issues (83%), musculoskeletal defects (83%), cardiac issues (65%), kidney issues (61%), and dysmorphic features (37%) (**Fig. 1A**). Of note, these summary-level rates are higher than what has been reported in the literature (Bonardi et al., 2021; McTague et al., 2018), potentially reflecting the more comprehensive nature of medical record review. When breaking down the respiratory symptoms, we observed that 78% of individuals were dependent on assisted breathing, with many having gone into respiratory failure (48%) and/or distress (41%), had excess fluids in their lungs (46%), experienced hypoxia (40%), and/or had structural lung defects (24%) (**Fig. 1B**). We also analyzed the cardiac issues and found that 40% of patients had abnormal electrocardiograms (ECGs) and about a third of these patients had heart murmurs (33%) and congenital structural defects (28%) of the heart (**Fig. 1C**). 13% of patients had systemic-pulmonary collaterals, formerly known as major aortopulmonary collateral arteries (MAPCAs) (Delaney et al., 2025) and 7% of patients had arrhythmias (**Fig. 1C**). Within the musculoskeletal defects, we found that about half of these patients had hypotonia (59%), hip deformities (54%) and joint contractures (43%) (**Fig. 1D**). A little over a third of patients had spine deformities (35%) which includes scoliosis of the spine, and compromised bone integrity (33%) (**Fig. 1D**). Within the reported kidney issues, almost half of these patients (43%) experienced urinary dysfunction including retention of urine and incontinence (**Fig. 1E**). 17% of patients experienced recurrent urinary tract infections (UTIs), 15% experienced renal stones, 13% experienced urinary obstructions, and 4% of patients were reported to have structural kidney defects (**Fig. 1E**). We also found that 22% of patients had dysmorphic features, 20% had plagiocephaly, 11% had foot deformities, and 4% had leg length inequality (**Fig. 1F**). After seizures, respiratory issues had the second highest prevalence within KCNT1 patients, with the vast majority dependent on an assisted breathing device. These results demonstrate the variety of multi-organ symptoms these patients experience and highlight the need to develop more targeted treatments by shedding light on the endogenous developmental role of KCNT1 across organ systems.

**Figure 1.**
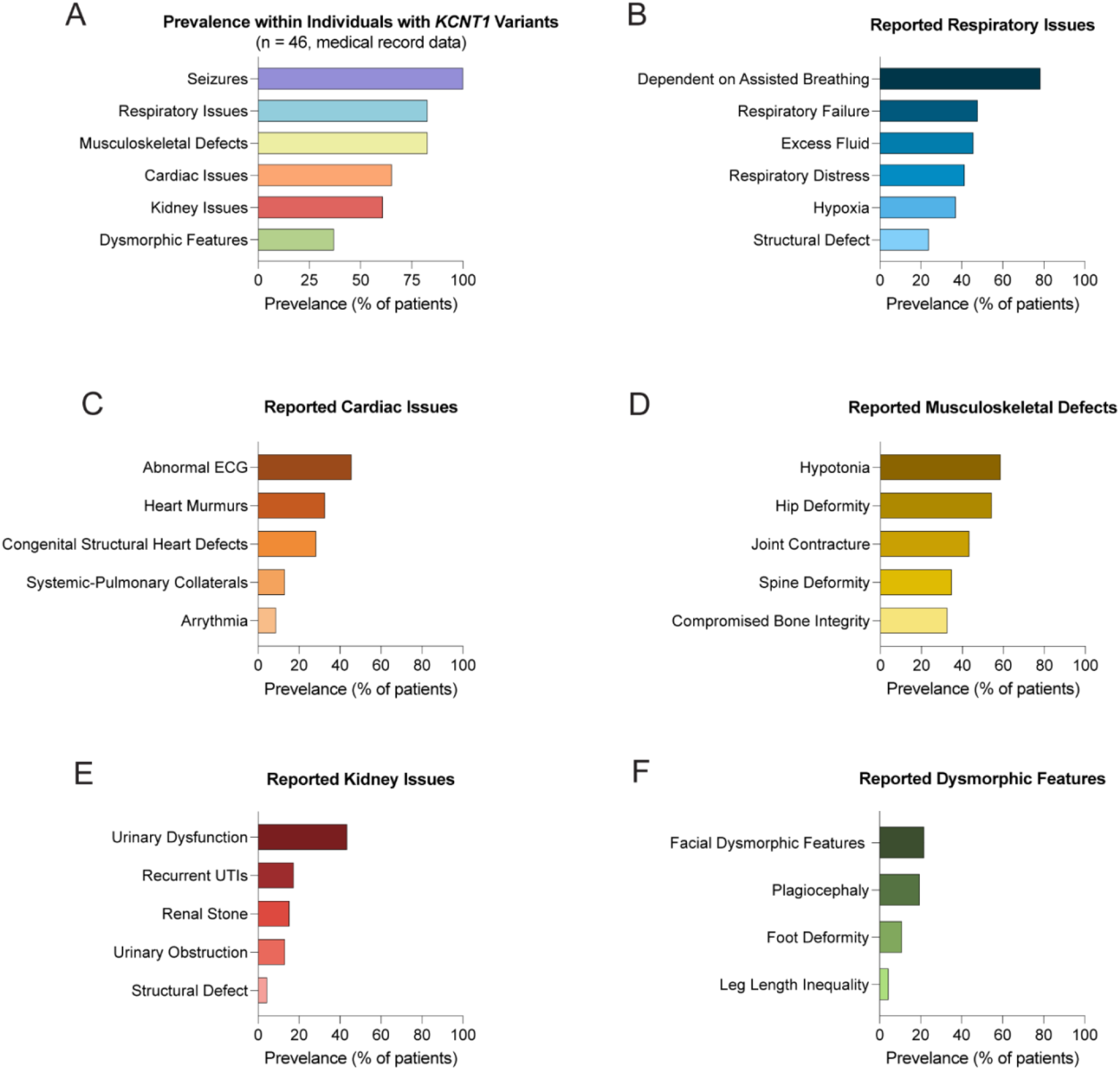
Prevalence of clinical features among individuals with pathogenic *KCNT1* variants. (A) Patient medical record data representing 46 patients with variants in *KCNT1* reveal prevalence of clinical symptoms affecting many organ systems. (B) Within the respiratory issues, 78% of patients were dependent on assisted breathing, 48% experienced respiratory failure, 46% had excess fluid in their lungs, 41% experienced respiratory distress, 40% experienced hypoxia, and 24% had structural defects. (C) Within the cardiac issues, 40% of patients had abnormal ECG, 33% had heart murmurs, 28% had congenital structural heart defects, 13% had systemic-pulmonary collaterals, and 7% had arrhythmias. (D) Within the musculoskeletal defects, 59% of patients had hypotonia, 54% had hip deformities, 43% had joint contractures, 35% had spine deformities, and 33% had compromised bone integrity. (E) Within the kidney issues, 43% of patients had urinary dysfunction, 17% had recurrent UTIs, 15% had renal stones, 13% had urinary obstruction, and 4% had structural defects. (F) Within the dysmorphic features, 22% of patients had facial dysmorphic features, 20% had plagiocephaly, 11% had foot deformities, and 4% had leg length inequality. See also Table S1.

### kcnt1 is broadly expressed in developing X. tropicalis embryos, including in epidermal multiciliated cells

*Xenopus tropicalis* is a diploid frog model amenable to genetic analysis and transcriptomics throughout embryogenesis (Grainger, 2012; Willsey et al., 2024); therefore first we sought to characterize the expression pattern of *kcnt1* over development in this vertebrate species to test the hypothesis that *kcnt1* is widely expressed across organ systems in developing embryos. Using whole-mount RNA *in situ* hybridization in this species, we observed that *kcnt1* is expressed early in development in the neural tube (which becomes the central nervous system) and in the neural crest (which becomes the facial structures and peripheral nervous system) at stage 20 (**Fig. 2A**). We observed *kcnt1* expression in the otic vesicle (which becomes the auditory system), pharyngeal arches (which become the facial structures), and multiciliated epidermis at stage 30 (**Fig. 2A**). In addition, *kcnt1* was also highly expressed in the brain, eye, heart and pronephros (developing kidney) at stage 35, all of which are highly ciliated organs (**Fig. 2A**). Similarly, *kcnt1* has been shown to be expressed in several regions of the brain, otic vesicle and olfactory organ in developing zebrafish embryos (Silic et al., 2021). These results are consistent with the hypothesis that KCNT1 has broad roles in embryonic development and are consistent with affected patients experiencing an early-onset, multi-organ clinical presentation when KCNT1 function is perturbed from birth.

**Figure 2.**
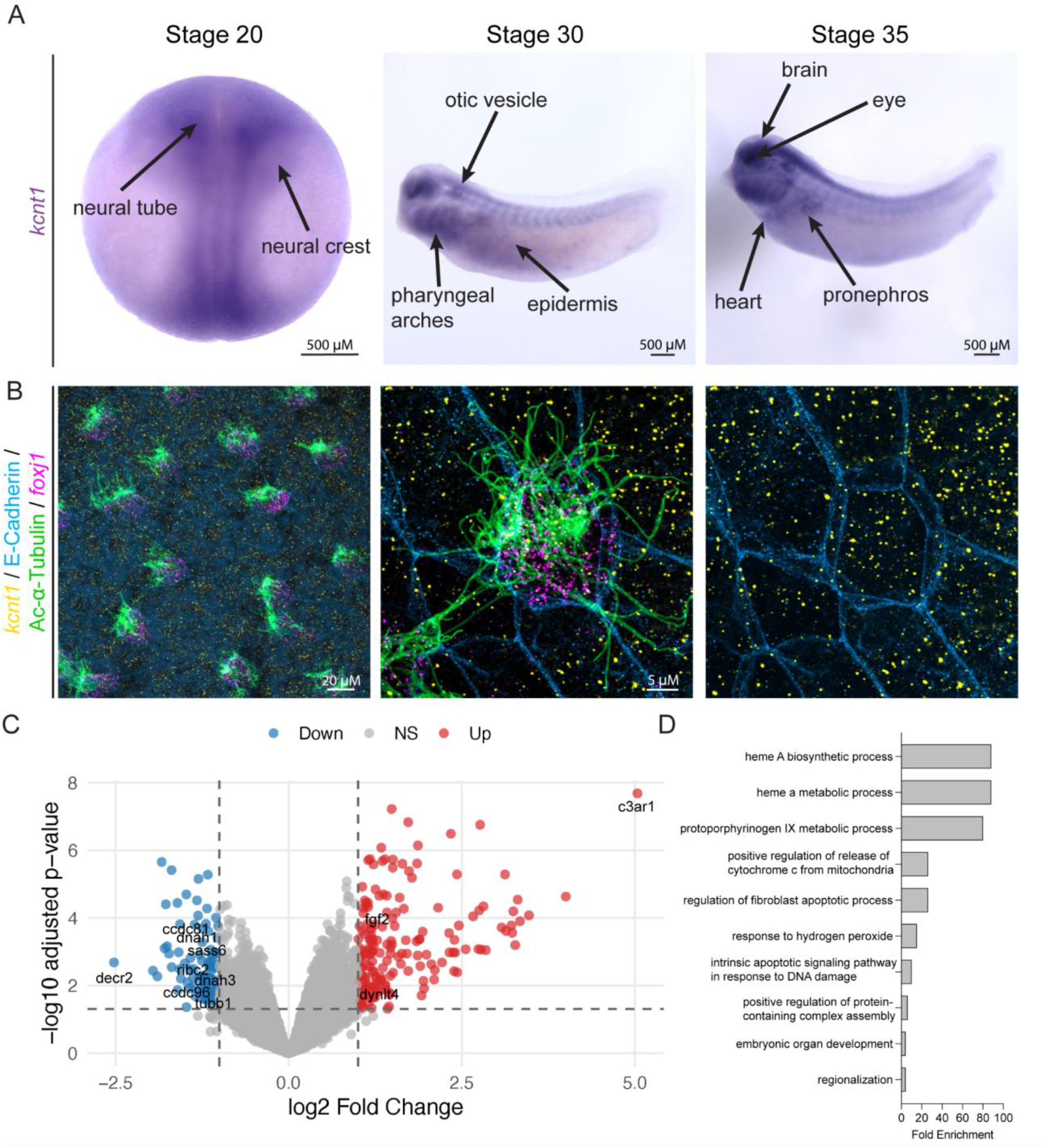
*kcnt1* is expressed in ciliated tissues, including the epidermis, during embryonic organ development. (A) *kcnt1* is expressed in ciliated embryonic tissues including the neural tube, epidermis, heart and pronephros in NF stages 20, 30, and 35 in *X. tropicalis* embryos. (B) *kcnt1* (yellow, HCR probe) is broadly expressed on the ciliated epidermis, labeled with cell boundary protein E-cadherin (blue, antibody staining), cilia protein acetylated alpha-tubulin (green, antibody staining), and MCC marker *foxj1* (magenta, HCR probe), in developing *X. tropicalis* embryos in NF stage 30. (C) Volcano plot of bulk RNA sequencing of embryos depleted of *kcnt1* with translation-blocking morpholino in *X. tropicalis* with select significant genes, sequenced at stage 33, compared to control morpholino-injected animals. (D) Top 10 biological processes organized by adjusted p value followed by fold enrichment using GO term analysis of differentially expressed genes in C. See also Table S2, S3.

Given the rates of severe respiratory issues and structural heart defects in this patient population, we wanted to specifically assay KCNT1 expression in relevant developing ciliated cell types. Motile cilia are critical for both respiratory clearance, through fluid flow generated by motile multiciliated cells, and for heart development, via motile cilia-generated fluid flow at the embryonic node (Bustamante-Marin and Ostrowski, 2017; Legendre et al., 2021; Mitchell et al., 2007; Shaikh Qureshi and Hentges, 2024). In *Xenopus*, the embryonic epidermis is a canonical model system for studying motile cilia form and function, particularly given its strong orthology to the human airway (Kulkarni et al., 2018; Rao et al., 2025; Sim et al., 2018; Ventrella et al., 2023; Walentek, 2021; Walentek et al., 2015; Werner et al., 2011). Therefore, to specifically query whether KCNT1 is expressed in motile multiciliated cells, we performed fluorescent RNA *in situ* hybridization chain reaction (Willsey, 2021) and observed *kcnt1* broadly expressed (yellow) across cell types in the developing multiciliated epidermis in *X. tropicalis* embryos (stage 30), including within multiciliated cells marked by *foxj1* expression (magenta) and acetylated-alpha Tubulin antibody staining (green) (**Fig. 2B**). Together, these results show that *kcnt1* is broadly expressed in embryonic ciliated tissues across developing organ systems, including the highly multiciliated epidermis in *Xenopus*, a common model system for the human airway. These results are consistent with a potential endogenous function for KCNT1 in embryonic ciliary biology that may be relevant to patient symptomatology.

### kcnt1 loss of function perturbs expression of key developmental signaling pathways

To investigate the function of *kcnt1* in the developing embryo, we depleted *X. tropicalis* embryos of Kcnt1 protein by injecting a translation-blocking morpholino in two blastomeres at the four-cell stage targeting the epidermis, and compared to control non-targeting morpholino-injected embryos. We performed bulk RNA sequencing on these embryos at stage 33 in triplicate and identified 253 differentially expressed genes ( *P* values ≤ 0.05, with 177 increased in expression and 76 decreased) (**Fig. 2C, Table S2**). 81 gene ontology terms within the “Biological Processes” category were significantly enriched (FDR-adjusted *P* values ≤ 0.05) (**Fig. 2D, Table S3**). Among the top 10 terms ranked by adjusted *P* value followed by fold enrichment included “embryonic organ development” and “regionalization” (**Fig. 2D**), consistent with a broad role for KCNT1 in embryonic development.

Among differentially expressed genes, the major developmental pathway regulator, fibroblast growth factor 2 (*fgf2*) was significantly misexpressed, and its receptor has been shown to promote ciliary growth and function (Kunova Bosakova et al., 2019; Nita et al., 2025; Yuan et al., 2019). In addition, genes encoding ciliary components like RIB43A domain with coiled-coils 2 (*ribc2*), tubulin beta class 1 (*tubb1*), spindle assembly abnormal protein 6 homolog (*sass6*) and coiled-coil domain containing 81 and 96 (*ccdc81* and *ccdc96*) were misexpressed following Kcnt1 depletion (**Fig. 2C, Table S2**). Similarly, known ciliary components were significantly misexpressed, including motile cilia motors including dynein axonemal heavy chain 1 and 3 (*dnah1, dnah3*) and dynein light chain Tctex-type 4 (*dynlt4*) (**Fig. 2C, Table S2**). Several of these genes (*ccdc96, ribc2*, and *dnah3*) are specifically required in motile cilia (Bazan et al., 2021; Kwon et al., 2023; Wang et al., 2024), suggesting that KCNT1 may be particularly relevant to motile cilia. Together, these results are consistent with KCNT1 having broad roles in embryonic development and potentially specifically within ciliary biology.

### kcnt1 is required for Xenopus multiciliated cell development

In *Xenopus*, the embryonic mucociliary epidermis is particularly amenable to rapid ciliary phenotyping because the multiciliated cells are abundant, regularly spaced throughout the surface of the embryo, and easily identified by antibody staining (Deblandre et al., 1999; Teerikorpi et al., 2025; Werner and Mitchell, 2013). Therefore, we used this system to specifically test whether loss of KCNT1 affects embryonic ciliary biology. To do this, we injected the same translation-blocking *kcnt1* morpholino or control morpholino, along with an H2B-GFP tracer, and targeted the embryonic mucociliary epithelium by one of four-cell stage injection. Injected embryos (marked by H2B-GFP) were stained with canonical cilia marker acetylated alpha-tubulin antibody and evaluated blinded to condition and categorized as having severe, moderate, mild or no ciliary defects (categorical examples in **Fig. SF1**). We observed that embryos depleted of *kcnt1* had significantly more severe cilia phenotypes, compared to embryos injected with control morpholino (**Fig. 3A-C**, magenta). Critically, the multiciliated cells seem specifically affected since the epidermis of these kcnt1-depleted embryos remained intact as assayed by F-actin phalloidin staining. (**Fig. 3A, 3C**, grey).

**Figure 3.**
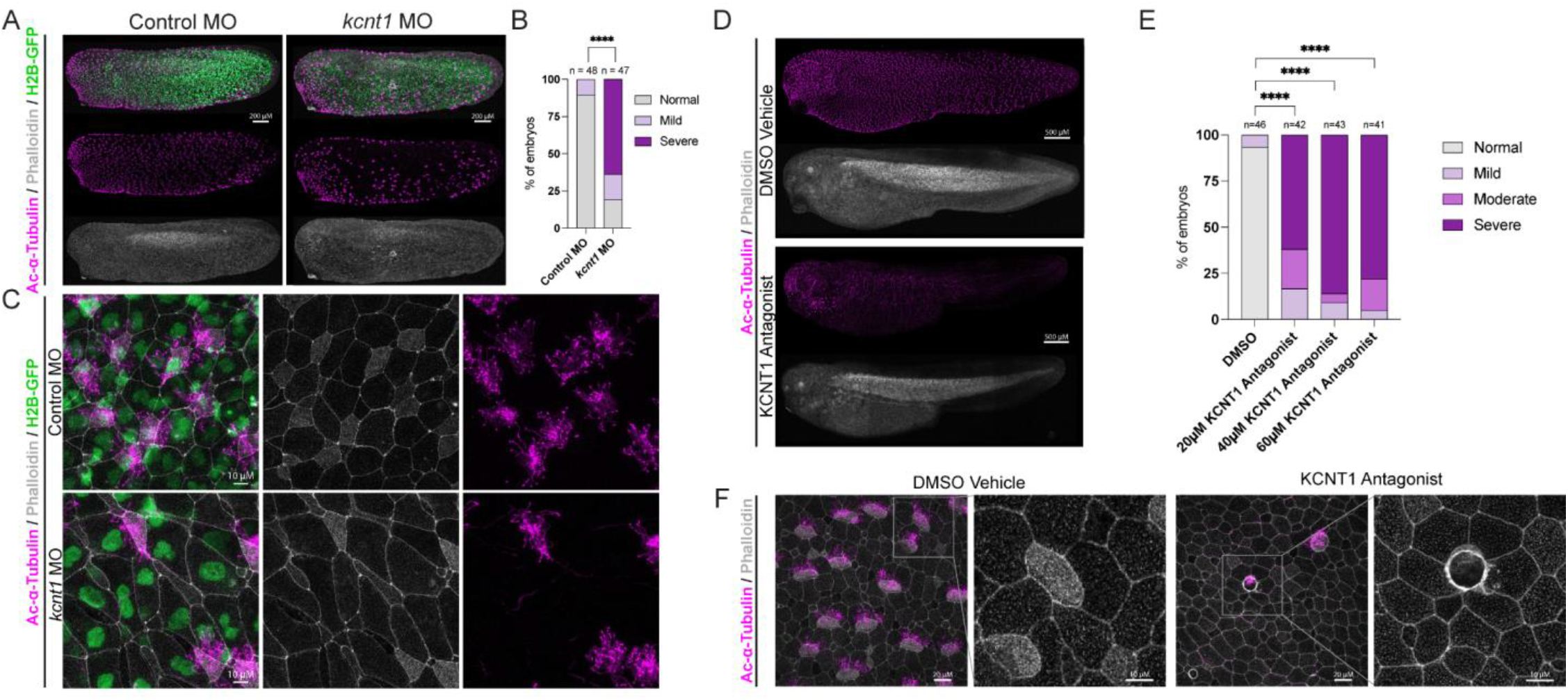
Depletion or inhibition of KCNT1 results in loss of motile cilia. (A) Depletion of *kcnt1* by a translation-blocking morpholino (MO) reduces motile cilia compared to control morpholino-injected NF stage 28 *X. tropicalis* stained for acetylated-α-tubulin (cilia, magenta) and phalloidin (actin, grey), and injected with tracer H2B-GFP (nucleus, green). Scale bars = 200 µm. (B) Quantification of data shown in A as a percentage of embryos with no cilia phenotype vs. mild, moderate or severe cilia phenotype. See Fig. SF1 for representative images of cilia phenotype classifications. Embryo images were randomized and phenotyped blinded to condition. Chi-Square test was used to calculate significance using raw counts of individual embryos. **** = p < 0.0001. (C) Epidermis of embryos from A. Scale bars = 10 µm. (D) *X. laevis* embryonic treatment with KCNT1 antagonist VU0606170 reduces motile cilia compared to DMSO vehicle treated from NF stage 7 to NF stage 38 *X. laevis*. Embryos were stained with acetylated-α-tubulin (cilia, magenta) and phalloidin (actin, grey). Scale bars = 500 µm. (E) Quantification of data shown in D as a percentage of embryos with no cilia phenotype vs. mild, moderate or severe cilia phenotype. Chi-Square test was used to calculate significance using raw counts of individual embryos. **** = p < 0.0001. (F) Epidermis of embryos from D. Scale bars = 20 µm followed by 10 µm. See also Fig. SF2 for treatment with an additional KCNT1 inhibitor, Compound 31 (Griffin et al., 2021), which also phenocopies these results.

To test the specificity of these findings, we attempted to pharmacologically phenocopy this KCNT1 inhibition. Previous work identified a KCNT1 inhibitor, VU0606170, that specifically reduced channel activity of cells overexpressing KCNT1 (reference sequence and patient variant sequence), but not KCNT2 (Spitznagel et al., 2020). In addition, this inhibitor was tested against a structurally similar compound and only VU0606170 reduced neuronal firing (Spitznagel et al., 2020). Therefore, we tested whether this KCNT1 inhibitor also causes a cilia phenotype in *Xenopus laevis*, similar to the Kcnt1 morpholino in *Xenopus tropicalis*. To do this, we treated *X. laevis* embryos with varying doses including 20 µM, 40 µM and 60 µM VU0606170 starting at stage 7, compared to DMSO-treated animals in parallel. Again, we were blinded to the condition and assessed ciliated cells on the epidermis of each embryo using acetylated alpha-tubulin antibody staining (magenta), categorizing embryos as having severe, moderate, mild, or no ciliary defects. We observed even at lower doses of VU0606170 (20 µM), there was still a very strong effect on ciliation with the vast majority of embryos having a severe ciliary phenotype (**Fig. 3D-F**, magenta). As with the morpholino, these embryos still had epidermal cells, as assayed by F-actin phalloidin staining (**Fig. 3D, 3F**, grey). We further validated these findings by treating embryos with another KCNT1 inhibitor, Compound 31, which has been shown to selectively target KCNT1 and reduce seizures in a GOF mouse model (Griffin et al., 2021). We treated embryos with 50 µM and 75 µM of Compound 31 starting at stage 7, compared DMSO animals in parallel and categorized embryos into severe, moderate, mild or no ciliary defects. 50 µM of Compound 31 resulted in more of a mild ciliary defect, while 75 µM resulted in the majority of embryos with either a moderate or severe ciliary defect (**Fig. SF2A-C**, magenta). Consistent with the morpholino and previous drug treatment, the epithelium of these embryos remained intact as assayed by F-actin phalloidin staining (**Fig. SF2A, SF2C**, grey). While imaging the phalloidin staining of VU0606170 treated embryos, we noticed that embryos treated with this inhibitor had more macropinocytotic events than usual (**Fig. 3F**). These are seen as actin ruffles on the plasma membrane, thought to engulf extracellular fluid and cargo into large vesicles, and often form in this tissue when it is under mechanical stress from crowding (Bresteau et al., 2025; Kay, 2021). Together these results indicate that KCNT1 is required during embryonic epidermis formation for multiciliated cell development and suggest that KCNT1 may play a role in mechanosensation.

### Kcnt1 interacts with Piezo to regulate ciliogenesis

Since macropinocytosis is often seen on the ciliated epidermis in response to mechanical stretch (Bresteau et al., 2025), we next wanted to probe its interaction with the mechanosensory signaling channel Piezo1. In *Xenopus*, Piezo signaling inhibition increases macropinocytotic events (Bresteau et al., 2025) and leads to loss of MCC specification and reduced *foxj1* expression (Narayanan et al., 2025), a master transcription factor that specifies MCC fate (Yu et al., 2008). In contrast, Piezo1 activation decreases macropinocytotic events (Bresteau et al., 2025) and promotes ciliation (Miyazaki et al., 2019). Since we observed that KCNT1 inhibition increased macropinocytotic events, we hypothesized that Kcnt1 signals through Piezo1 to regulate MCC development, consistent with previous reports of potassium-mediated regulation of Piezo1 (Mitchell et al., 2025). To test this hypothesis, we co-treated *X. laevis* embryos with 20 µM Kcnt1 antagonist, VU0606170, and 2.5 µM Piezo antagonist, GsMTx-4, starting at stage 7 and phenotyped epidermal multiciliated cells of the embryonic epidermis at stage 38 by acetylated alpha-tubulin antibody staining, blinded to condition. We observed that co-treatment of the KCNT1 and Piezo inhibitors caused almost complete loss of multiciliated cells, more severe than the KCNT1 antagonist treatment alone (**Fig. 4A-B**, magenta). Conversely, co-treatment of 20 µM KCNT1 antagonist with 110 µM of Piezo1 agonist, Yoda1, starting at stage 7 in *X. laevis* partially rescued the ciliogenesis defect observed with the KCNT1 antagonist alone (**Fig. 4C-D**, magenta). As before, these embryos still had epidermal cells, as assayed by F-actin phalloidin stain (**Fig. 4A, 4C**, grey). Together, our results support the hypothesis that KCNT1 functionally interacts with Piezo, which is known to control *FOXJ1* expression and multiciliated cell development (Narayanan et al., 2025). This is consistent with previous work in mammalian systems demonstrating the impact of potassium ion flux upstream of Piezo signaling (Mitchell et al., 2025). Together, this work suggests a broad role for KCNT1 in embryonic development, across organ systems, and specifically in multiciliated cell development. These results are consistent with the patient presentation and offer new opportunities for therapeutic design, particularly in the case of the severe respiratory issues seen in affected individuals.

**Figure 4.**
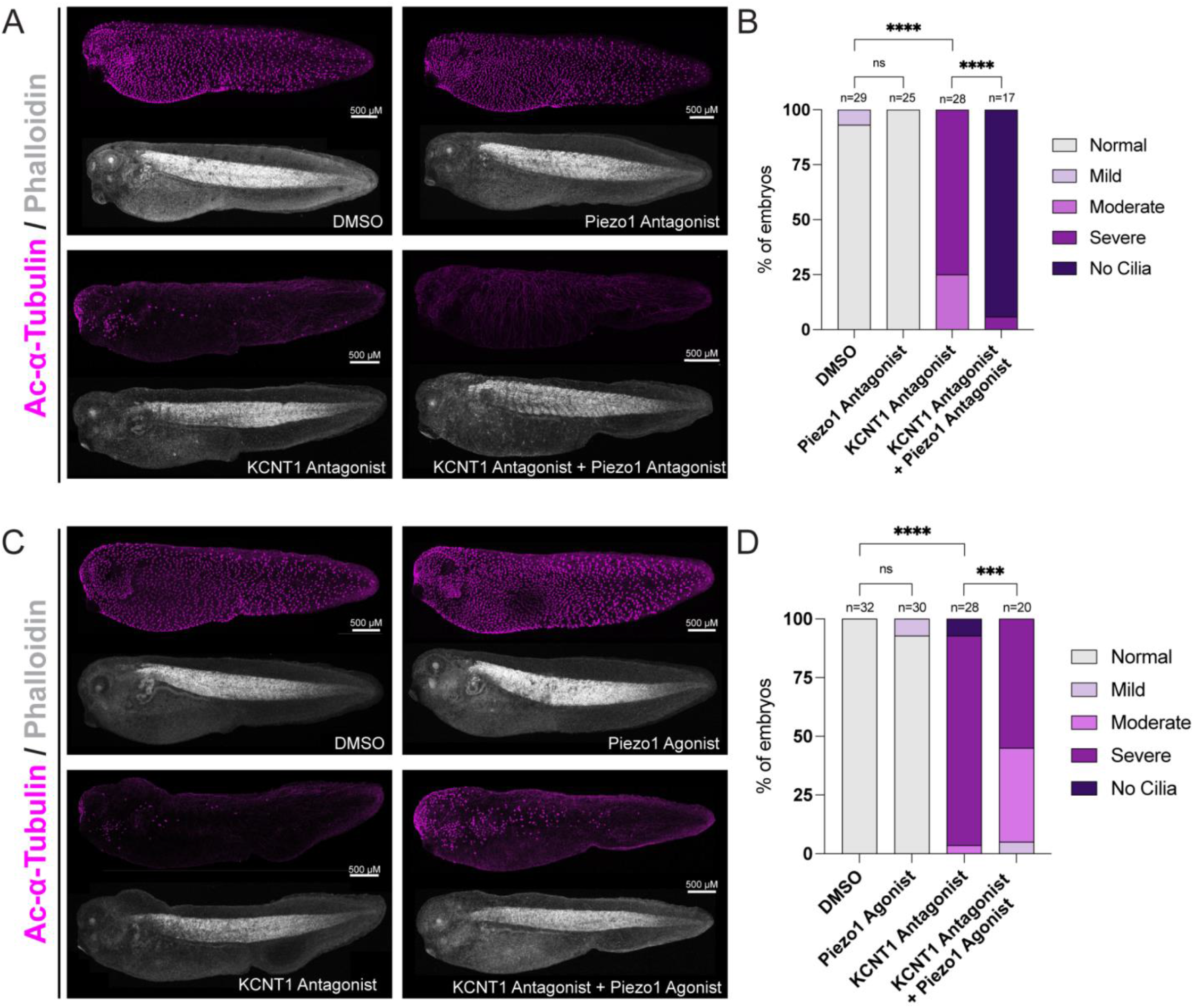
Inhibition of KCNT1 is exacerbated by Piezo signaling inhibition and can be partially rescued by Piezo1 activation. (A) Inhibition of KCNT1 and Piezo signaling with KCNT1 antagonist, VU0606170 (20 µM), and Piezo signaling antagonist, GsMTx4 (5 µM), further exacerbates loss of motile cilia compared to KCNT1 antagonist (20 µM) alone. *X. laevis* embryos were treated from NF stage 6 to 38 and stained with acetylated-α-tubulin (cilia, magenta) and phalloidin (actin, grey). Embryo images were randomized and scored blinded to condition. Scale bars = 500 µm. (B) Quantification of data shown in A as a percentage of embryos with no cilia phenotype vs. mild, moderate or severe cilia phenotype. Fisher’s exact test was used to calculate significance using raw counts of individual embryos. **** = p < 0.0001, ns = no statistically significant difference. (C) Partial rescue of loss of cilia phenotype from KCNT1 inhibition (20 µM VU0606170) by co-treatment with a Piezo1 agonist, Yoda1 (110 µM), compared to KCNT1 antagonist alone (20 µM). Scale bars = 500 µm. (D) Quantification of data shown in C as a percentage of embryos with no cilia phenotype vs. mild, moderate or severe cilia phenotype. Fisher’s exact test was used to calculate significance using raw counts of individual embryos. **** = p < 0.0001, *** = 0.0009, ns = no statistically significant difference.

## Discussion

Pathogenic variants in *KCNT1* are classically associated with severe early-onset epileptic encephalopathies, yet the substantial extra-neurological burden observed in affected individuals has remained insufficiently explained (Barcia et al., 2012; Bonardi et al., 2021; McTague et al., 2018). By integrating clinical cohort analysis with embryological studies in *Xenopus*, our findings reveal an additional, previously underappreciated role for KCNT1 in development, expanding its function beyond its well-established role in neuronal excitability. The convergence of multi-organ clinical involvement with embryonic expression in ciliated tissues suggests that KCNT1 participates in conserved epithelial and developmental programs relevant to multiple organ systems. Our clinical review reinforces that KCNT1-associated disease frequently involves systems not typically emphasized in genetic epilepsies, including congenital heart defects, respiratory dependence, urinary dysfunction, and spinal abnormalities. Although retrospective and limited by cohort size, the recurring pattern of involvement across these systems argues for shared developmental mechanisms rather than unrelated comorbidities. Many of the affected organs either contain motile cilia or depend on cilia-mediated signaling during development, providing a biologically coherent framework for further investigation.

Consistent with this clinical signal, *kcnt1* transcripts were detected in multiple developing ciliated tissues in *Xenopus*, including the mucociliary epidermis, pronephros, heart, and neural regions. Functional perturbation of Kcnt1 produced robust abnormalities in multiciliated cell biology within the mucociliary epidermis, a well-established proxy for the human airway epithelium (Walentek, 2021; Walentek and Quigley, 2017). Importantly, our data do not distinguish whether KCNT1 primarily influences early cell fate decisions, later ciliogenesis processes, or both. Rather, the findings place KCNT1 upstream of normal multiciliated cell development and function in this tissue context. Given the prominent respiratory morbidity reported in patients, these results provide a plausible mechanistic link that warrants deeper investigation in mammalian airway models and models of *KCNT1* patient-derived variants.

The interaction between KCNT1 and Piezo signaling further suggests that ion channel cross-talk may be an important regulatory feature of multiciliated epithelia. Pharmacologic inhibition of Piezo signaling enhanced the KCNT1-associated phenotype, whereas Piezo1 activation partially mitigated it. These data support a model in which KCNT1 and Piezo1 function within a shared or convergent pathway governing epithelial differentiation programs. One possibility is that KCNT1-dependent regulation of membrane potential or ionic flux modulates mechanosensitive signaling thresholds, thereby influencing downstream transcriptional networks. However, the precise molecular relationship between these channels remains to be defined and will require direct electrophysiological and pathway-level studies.

Several important limitations should be considered. First, our experimental paradigm models KCNT1 loss of function, whereas the vast majority of patient variants are thought to confer gain-of-function channel activity. Although developmental systems often exhibit nonlinear sensitivity to ion channel dosage and activity, caution is warranted in extrapolating mechanistic directionality from this current model to human disease. Future studies using patient-specific gain-of-function variants, including knock-in and overexpression approaches, will be essential to determine whether altered channel activity perturbs multiciliated tissues through shared or distinct mechanisms. Second, the present work focuses on a single multiciliated epithelial population: the *Xenopus* mucociliary epidermis. While this tissue provides powerful experimental access and strong relevance to airway biology (Walentek, 2021; Walentek and Quigley, 2017), multiciliated cells are heterogeneous across organs, and primary cilia, which play especially critical roles in neuronal development and signaling (Atkins et al., 2025; Zhang et al., 2025), were not directly examined here. Given the profound neurological manifestations of KCNT1 - associated disorders, determining whether KCNT1 influences primary cilia structure or signaling in neural contexts is an important next step. Extending these studies to neuronal systems, ventricular zone epithelia, and other primary cilia-dependent tissues will be necessary to fully define the scope of KCNT1 developmental ciliary functions.

In summary, our findings expand the functional landscape of KCNT1 beyond its established role in neuronal excitability and support a broader contribution to embryonic epithelial biology. By linking patient phenotypes with conserved developmental processes and identifying interaction with the pharmacologically tractable Piezo pathway, this work opens new directions for understanding the multisystem manifestations of KCNT1-associated disorders. More broadly, these results contribute to a growing recognition in developmental biology that ion channels traditionally studied in excitable cells can also have instructive roles in tissue morphogenesis and epithelial differentiation.

## Acknowledgments

We thank the KCNT1 community for participating in research. We thank: Citizen Health for access to their patient medical record data; Ali Rosenberg and Justin West for helpful conversations; Jeremy Reiter and Mark von Zastrow for helpful suggestions and feedback; David Breckenridge for suggesting the use of an additional KCNT1 inhibitor; Nolan Wong and UCSF LARC for animal care; Ethel Bader, Catherine Nguyen, Juan Arbelaez, Jeanselle Dea, James Schmidt, and Milagritos Alva for laboratory maintenance and support; and Ashley Clement, Gigi Paras, Sonia Lopez, and Linda Chow for administrative support. This work would not be possible without daily reference to the Xenopus community resource Xenbase (RRID:SCR_003280) and expertise and frog resources from the National Xenopus Resource (RRID:SCR_013731). This work was supported by an investigator award from Biohub - San Francisco (to HRW).

## Author Contributions

Conceptualization: HRW. Methodology: AKC, KEM. Formal Analysis: AKC, MNT, ATE. Investigation: AKC, ATE. Resources: HRW. Writing - Original Draft: AKC. Writing - Review & Editing: HRW, AKC. Visualization: AKC, HRW. Supervision: HRW. Project Administration: HRW. Funding Acquisition: HRW.

## Materials and Methods

### Citizen Health patient data analysis

Citizen Health medical record data was determined to be non-human subjects research by UCSF IRB office, IRB#23-39-079. 8/46 patients had a pathogenic or likely pathogenic variant in another gene beyond *KCNT1*, but were still included in these analyses. 1/46 patients had a *KCNT1* variant classified as benign, but was still included in these analyses. Data analysis was performed in R studio using an established pipeline from Kostynavoksya 2025 (Kostyanovskaya et al., 2025) and visualization was performed in Prism software. Details of which symptoms fell into each category can be found in Supplementary Table 1.

### Xenopus husbandry and animal care

Both *Xenopus laevis* and *Xenopus tropicalis* were used in this study. *In situ* hybridization and morpholino experiments were done in *X. tropicalis*. Drug treatments were performed in *X. laevis*. All frog care was performed according to UCSF IACUC protocol #AN199587-00A. All *Xenopus* stages were based on the Normal Table of *Xenopus Laevis* (Daudin) (Hubrecht Laboratory (UTRECHT) et al., 1956). Xenbase (RRID:SCR_003280) was used for anatomical resources, phenotypes, and genetic references (Fisher et al., 2023, 2022; Segerdell et al., 2013). Wildtype frogs were supplied by the National Xenopus Resource Center (RRID: SCR_013731) (Pearl et al., 2012).

### Xenopus whole mount RNA in situ hybridization

*Xenopus tropicalis kcnt1* probe plasmid (*Xenopus* Gene Collection Clone 7653330) was a kind gift from Dr. Richard Harland (UC Berkeley). Digoxigenin-11-UTP-labeled antisense RNA probe was synthesized from this plasmid by standard protocol (Sive, 2000) using SalI restriction enzyme and T7 polymerase. Whole mount RNA *in situ* hybridization was performed on *X. tropicalis* stage 20, 30 and 35 embryos according to Willsey 2021 (Willsey, 2021) with the omission of the proteinase K step. Images were taken on Zeiss AxioZoom.V16 with a 1x objective and extended depth of focus for full-embryo imaging.

For hybridization chain reaction (HCR), embryos were stained according to Willsey 2021 (Willsey, 2021), with an additional step of heating probes to 95 °C for 90 seconds and cooling to room temperature before use. *kcnt1* and *foxj1* probes were custom ordered through Molecular Instruments and designed to target the *X. tropicalis* genes. Embryos were imaged on a Zeiss 980 LSM with a 63x oil objective, Airyscan processed and Maximum Intensity Projection performed in Zen.

### *Xenopus* Bulk RNA sequencing data analysis

*Xenopus tropicalis* embryos were injected at the 4 cell stage into 2 blastomeres targeting the epidermis, and injected with 5.1ng each of *kcnt1* morpholino (details in “*Xenopus* gene perturbations”) and 100 pg of tracer H2B-GFP mRNA. Each replicate consisted of 3 embryos collected at NF stage 33 combined and lysed using the RNeasy kit for RNA purification (Qiagen 74104). Samples were submitted to Plasmidsaurus for bulk RNA sequencing. We used the aligned and sample-level feature counts provided by Plasmidsaurus for analysis using DESeq2 (version 1.48.2) in R (version 4.5.1). Plasmidsaurus provided 3’ end counting for transcriptome-wide differential gene expression data and utilized the Illumina platform to provide ∼10M deduplicated 3’ end counting reads (20M raw reads) from 300 ng purified RNA. We first filtered lowly expressed genes, defined by requiring at least two samples to express more than 10 transcript counts per sample, then performed differential expression analysis, using the DESeq function, modeling on KCNT1 knockdown status (default parameters). We defined significant differentially expressed genes as those with false discovery rate (FDR)-adjusted *P* value ≤ 0.05. We used the clusterProfiler package (version 4.16.0) for gene ontology over-representation analysis, converting differentially expressed genes to human Entrez ID as input, and querying enriched gene ontology terms in all ontologies: molecular function (MF), biological process (BP), and cellular component (CC).

### Xenopus immunofluorescence

The following primary antibodies were used: E-cadherin (BD Bioscience 610181, 1:100) and Acetylated-α-Tubulin (Sigma T6793, 1:2000). The following secondary antibodies were used at 1:1000: anti-mouse-555 (Thermo Fisher Scientific, A32727), anti-mouse-647 (Thermo Fisher Scientific, A32728). All samples were fixed in 4% paraformaldehyde for 45 minutes then underwent standard immunofluorescence staining (Willsey, 2021) with the omission of bleaching. Phalloidin (Life Technologies A22287, 1:400) were incubated along with the secondary antibodies.

### Xenopus gene perturbations

For *Xenopus tropicalis* knockdown experiments, morpholinos were purchased from Gene Tools, and their sequences are: *kcnt1* (5’-CGGGACTCTCCTGTTTGGCTATAAA-3’) and standard control (5’- CCTCTTACCTCAGTTACAATTTATA-3’). For both *kcnt1* and the standard control morpholino, 5.1 ng of morpholino was injected into 1 blastomere of 4-cell stage *X. tropicalis* embryos, along with 100 pg of tracer H2B-GFP mRNA. Centrin-CFP, which labels MCC basal bodies, was initially used as a tracer; however, Centrin-CFP was not observed in *kcnt1* morpholino injected embryos, whereas control morpholino injected embryos exhibited numerous Centrin-CFP–positive multiciliated cells. Therefore, we switched to using H2B-GFP, which was robustly maintained in both conditions.

### *Xenopus* drug treatments

KCNT1 antagonist (VU0606170, ProbeChem PC-73240), additional KCNT1 antagonist (Compound 31, Enamine EN300-27781598), Piezo1 antagonist (GsMTx4, MedChemExpress HY-P1410), and Piezo1 agonist (Yoda1, Sigma SML1558) were resuspended in DMSO at 10 mM, 10 mM, 5 mM, and 10 mM, respectively. *Xenopus laevis* embryos were treated with drug or an equal volume of DMSO, diluted in 1/3 MR, at NF stage 7 and fixed at NF stage 38 for immunofluorescence staining. Dead embryos were removed as soon as they were discovered.

### Imaging and image analysis

All images were acquired on a Zeiss LSM 980 confocal microscope with either 5x air or 63x oil objective. Images were acquired in confocal mode. Images were processed in ImageJ (v2.0.0). Single plane images of embryos from all conditions were compiled into one folder and cilia phenotypes were scored using the Blind Analysis Tools from Image J, refer to Supplementary Figure 1 for representative images. Max intensity projections of representative embryos were taken with 5x air objective with z-stacks.

**Supplementary Figure 1.**
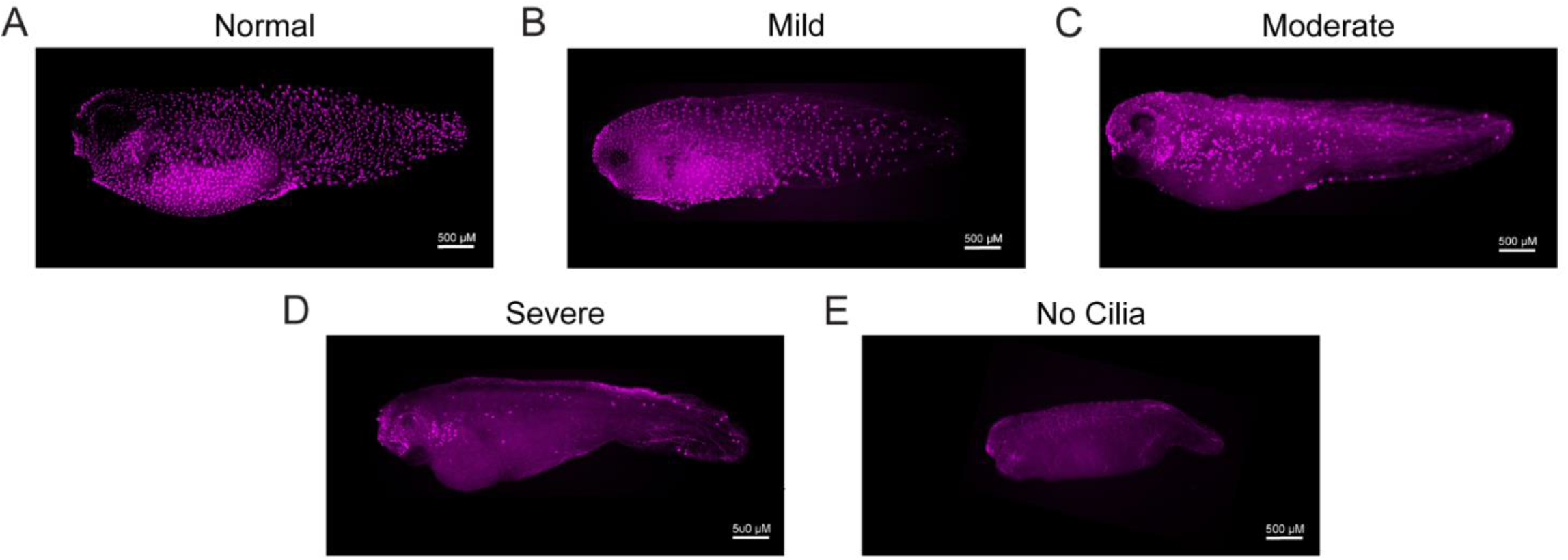
Representative images of cilia phenotyping. (A) Example of ciliary phenotype “Normal.” (B) Example of ciliary phenotype “Mild.” (C) Example of ciliary phenotype “Moderate.” (D) Example of ciliary phenotype “Severe.” (E) Example of ciliary phenotype “No cilia.”

**Supplementary Figure 2.**
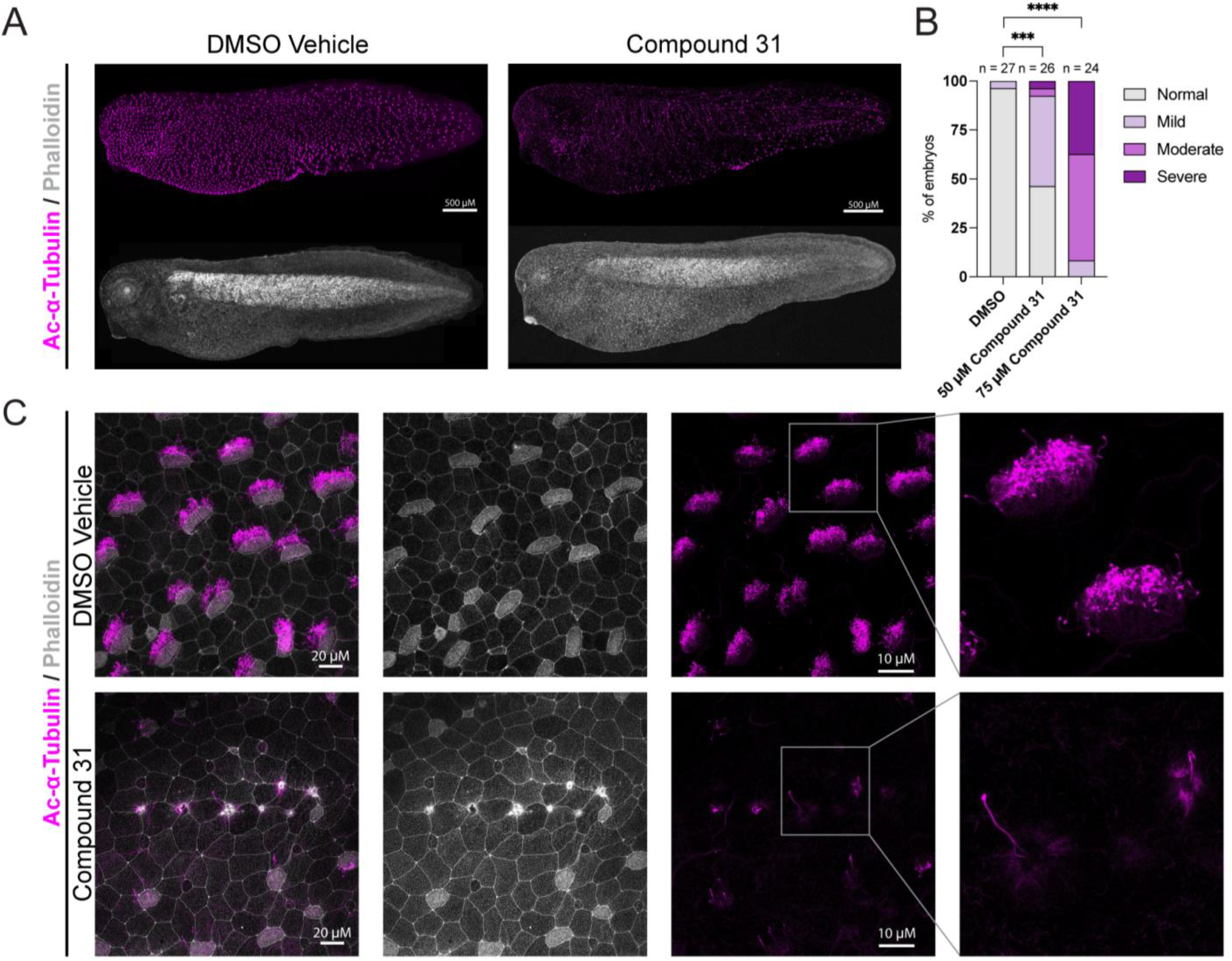
Inhibition of KCNT1 through Compound 31 results in loss of motile cilia. (A) Inhibition of KCNT1 with KCNT1 antagonist, Compound 31 (50 or 75 µM), reduces motile cilia compared to DMSO vehicle treated embryos, treated from NF stage 7 to NF stage 38 in *X. laevis*, stained with acetylated-α- tubulin (cilia, magenta) and phalloidin (actin, grey). Embryo images were randomized and scored blinded to condition. Scale bars = 500 µm. (B) Quantification of data shown in A as a percentage of embryos with no cilia phenotype vs. mild, moderate or severe cilia phenotype. Chi-Square test was used to calculate significance using raw counts of individual embryos. **** = p < 0.0001, *** = 0.0009. (C) Epidermis of embryos from A. Scale bars = 20 µm followed by 10 µm.

## Supplemental Table Legends

**Table S1**. Summary table categorizing symptoms from Citizen Health medical record review of *KCNT1* patients into respiratory, cardiac, musculoskeletal, renal, and dysmorphic groups, including counts for each category. See also Fig. 1.

**Table S2**. Spreadsheet containing results from bulk RNA sequencing comparing KCNT1 morpholino–injected embryos to control embryos. The first tab lists all detected genes with corresponding differential expression statistics, including gene name, expression values, log2 fold change, and associated p-values and adjusted p-values. The second tab includes only significantly differentially expressed genes with cutoff based on false discovery rate (FDR)-adjusted *P* value ≤ 0.05. See also Fig. 2.

**Table S3**. Spreadsheet containing results from Gene Ontology (GO) analyses on bulk RNA sequencing data from KCNT1 morpholino–injected embryos compared to control embryos. This sheet includes GO terms generated from differentially expressed genes. See also Fig. 2 and Table S2.

